# Viral dynamics and immune responses to foot-and-mouth disease virus in African buffalo *(Syncerus caffer)*

**DOI:** 10.1101/2021.11.24.469808

**Authors:** Eva Perez-Martin, Brianna Beechler, Katherine Scott, Lin-Mari de Klerk-Lorist, Fuquan Zhang, Georgina Limon, Brian Dugovich, Simon Gubbins, Arista Botha, Nicholas Juleff, Robyn Hetem, Louis van Schalkwyk, Francois F. Maree, Anna Jolles, Bryan Charleston

## Abstract

Foot-and-mouth disease (FMD) is one of the most important livestock diseases restricting international trade. While it is clear that African buffalo (*Syncerus caffer*) act as the main wildlife reservoir, viral and immune response dynamics during FMD virus acute infection have not been described before in this species. We used experimental needle inoculation and contact infections with three Southern African Territories serotypes to assess clinical, virological and immunological dynamics for thirty days post infection. Clinical FMD in the needle inoculated buffaloes was mild and characterised by pyrexia. Despite the absence of generalised vesicles, all contact animals were readily infected with their respective serotypes within the first 2-9 days after being mixed with needle challenged buffaloes. Irrespective of the route of infection or serotype there were positive associations between the viral loads in blood and the induction of host innate pro-inflammatory cytokines and acute phase proteins. Viral loads in blood and tonsils were tightly correlated during the acute phase of the infection, however, viraemia significantly declined after a peak at 4 days post infection (dpi), which correlated with the presence of detectable neutralising antibodies. In contrast, infectious virus was isolated in the tonsils until the last sampling point (30 dpi) in most animals. The pattern of virus detection in serum and tonsil swabs was similar for all three serotypes in the direct challenged and contact challenged animals.

We have demonstrated for the first time, that African buffalo are indeed systemically affected by FMD virus and clinical FMD in buffalo is characterized by a transient pyrexia. Despite the lack of FMD lesions, infection of African buffalo was characterised by high viral loads in blood and oropharynx, rapid and strong host innate and adaptive immune responses and high transmissibility.

## 1. Introduction

Foot and mouth disease (FMD) is an acute vesicular viral disease of domesticated and wild *Artiodactyla* characterized as highly contagious with a very short incubation period. In the acute stages of disease in FMD susceptible livestock, clinical signs include fever, blister-like lesions followed by erosions on the tongue, mouth, snout and feet [1]. FMD is one of the most important livestock diseases that is endemic in Africa and causes serious socio-economic impact in the livestock industry and inhibits international trade [2].

FMD virus (FMDV) is a small (30nm in diameter) roughly spherical, non-enveloped positive-sense single-stranded RNA picornavirus of the genus *Aphthovirus.* Given its serological diversity, FMDV is classified into 7 serotypes: A, O, Asia1 and C (also named Eurasian serotypes) and the Southern African Territories (SAT) 1, 2 and 3 with varying global distribution and causing indistinguishable disease [3]

FMDV is mainly transmitted directly from infected animals in close contact with naïve animals during acute infection. FMDV has a very high rate of transmission and R_0_ values during early stages of the disease, are considered to be 21-88 for cattle, 1-14 for sheep [4–6] and very recently, the R_0_ estimated for African buffalo was 5-15.8 [7]. In cattle, the onset of clinical signs occurs 3-4 days after infection and transmission occurs on average, 0.5 days after the appearance of clinical signs [8] when very high titres of virus are found in the damaged epithelium due to vesicle formation and vesicular fluid [9]. In contrast to cattle, African buffalo develop a sub-clinical or inapparent infection after being experimentally infected with high doses of the three SAT serotypes, while the same virus strains in young Nguni cattle caused fatal and acute FMD [10]. In cattle, the oropharyngeal mucosa is the primary site of replication after natural infection, with subsequent dissemination to the lungs followed by viraemia of about 3-5 days of duration [11]. The major mechanism of controlling FMDV infection is the induction of neutralizing antibodies which are detected as soon as 4 days post-infection (dpi), peak at 14 dpi and are maintained for very long periods of time (years) [1]. The humoral immune response induced by infection or vaccination, protects the animal against FMD but does not consistently prevent replication in the oropharynx and establishment of persistent infection or the carrier status [12].

An innate non-specific immune response based on type I and III IFN has been described to play a role in the early protective response against FMDV in pigs and cattle [13–15]. In fact, even though FMDV has developed mechanisms to antagonize the IFN response *in vitro* [16] type-I/III IFN is readily detected in serum after FMDV infection in cattle, pigs, mice and African buffalo [10, 17].

Enhanced production of acute-phase proteins (APPs), haptoglobin and serum amyloid A (SAA) in serum, have been described in cattle during acute infection with FMDV [18]. Interestingly, detection of APP has been used as an indicator of a range of infectious diseases to monitor progression of disease, as a marker to assess animal health and welfare at farms or slaughterhouses, antibiotic treatment efficacy and recently as a biomarker of other infections in African buffalo[19].

Most African buffalo in sub-Saharan Africa are endemically infected with all three SAT serotypes [20–22] and are also considered the main, and for some authors, the sole FMDV reservoir [23] as they may become persistently infected for many years [24, 25]. Controlling transboundary diseases such as FMD is critical to significantly improve livestock productivity in endemic regions and allow international trade in livestock products [26]. FMD control in sub-Saharan Africa provides unique challenges because the SAT serotypes are maintained in wildlife and act as a source of infection for livestock [27, 28]. Therefore, an important element of FMD control in livestock in Africa is understanding the pathogenesis and transmissibility in African buffalo. SAT2 is the most widely distributed serotype and is also the serotype most often associated with outbreaks in livestock and wildlife, followed by SAT1 and then SAT3 [29–31]. However, in contrast to cattle, little information is known about the viral dynamics, shedding, transmission rates, and host-immune responses during the acute infection in African buffalo.

Therefore, the aim of this study was to fill the knowledge gaps of FMDV infection dynamics and immune-pathogenesis in African buffalo following needle or direct contact infection with three SAT FMDV serotypes. Parameters such as viraemia, viral shedding, clinical outcome and fever, as well as the systemic levels of APP in serum and innate and adaptive immune responses were analysed. Despite the lack of visible clinical signs, infected buffaloes show high body temperature, high virus titres in blood and nasopharynx, and readily transmit the virus to naïve buffalo.

## Materials and methods

### Experimental design and sampling

Twenty-four African buffalo (*Syncerus caffer)* were donated by the Hluhluwe-Imfolozi Game Reserve, South Africa; confirmed free from antibodies to FMDV by the OIE Regional Reference Laboratory (ARC-OVI) and transferred to experimental animal facilities at Skukuza, State Veterinary Services (SVS), Kruger National Park (KNP). Animals were allowed one month for acclimatisation and daily monitoring of the health was performed throughout the experiment. Experimental protocols were approved by of the Department of Agriculture, Forestry and Fisheries (DAFF) (Section 20: 12/11/1/8/3/) and the SANParks Animal Ethical Committee (N013-12). Animals were sedated with etorphine hydrochloride and xylazine during experimental procedures and sample collection.

The 24 buffalo, 12 female and 12 male, aging between 10 and 24 months were randomly divided into six identical groups (four animals each). Animals in three groups were subepithelially challenged with either SAT1, SAT2 or SAT3 FMDV, at a dose of 2.5x 10^5^ TCID_50_ in the tongue. These groups will be referred in the manuscript as “needle infected” (NI) animals. Two days after the challenge, the remaining three groups of four naïve buffalo were mixed with each of the three inoculated groups; these animals are referred as “contact animals”. FMDV Infection dynamics was studied in the buffalo following needle and natural exposure of each of the SAT viruses during the acute phase for 30 days.

Buffalo were monitored for the presence of FMD clinical signs and sampled on days 0 (day of the needle infection), 2, 4, 6, 8, 11, 14 and 30 day post-infection (dpi). Sample collection included blood, oropharyngeal scraping (probang), nasal and tonsil swabs. Whole blood samples collected from the jugular vein were centrifuged to extract serum to measure pro-inflammatory cytokines (type I/III IFN, IFNγ, TNFα) and APP by ELISA; and specific humoral immune response measurement by virus neutralization test (VNT) and ELISA. Blood samples were also collected in EDTA for leucocyte counts immediately after collection on a Coulter T-890 (Beckman). EDTA blood, probang and tonsil swab samples were collected for the detection of FMDV by qRT-PCR and virus isolation. Probang samples were obtained by gentle erosion of the oropharyngeal epithelium with the probang cup [32]. Epithelium was resuspended in 3ml of probang buffer (Eagles-hepes supplemented with penicillin/streptomycin (Sigma)) and snap frozen in liquid nitrogen. Left and right palatine tonsils were swabbed individually with nylon brushes (Cytotak™ Transwab, Medical Wire), dipped in criovials containing 0.5 ml of probang buffer and snap frozen in liquid nitrogen[10]. Cotton nasal swabs (Salivette^R^) were soaked in 0.5ml of PBS and introduced into both nostrils to collect nasal fluid. Swabs were then centrifuged, the liquid collected and aliquoted. All samples were stored at −80C until processing.

### Viruses and cell lines

Virus isolates used for animal challenge were SAT1/KNP/196/91, SAT2/KNP/19/89 and SAT3/KNP/10/90 with accession numbers KR108948, KR108949 and KR108950, respectively. These viruses are originally from buffalo in KNP, isolated in primary porcine kidney cells (PK) and propagated in IB-RS-2 porcine cell line [10].

IB-RS-2 cells were also used for the virus neutralization assay. ZZR-127 goat epithelial cells were used for virus isolation from tonsil swabs and probang samples and sera [33]. MDBK-t2 cells (Madin-Darby bovine kidney) cells transfected with a plasmid expressing the human MxA promoter driving a chloramphenicol acetyltransferase (CAT) cDNA were used for the antiviral assay to detect Type I/III IFNs [34]. Cell lines were maintained in minimal essential medium (MEM) supplemented with nutrient F-12 (ZZR-127 cells), hepes, L-glutamine, 10% foetal calf serum and antibiotics (penicillin 100 U/ml and penicillin 100ug/ml). MDBK-t2 cells were also supplemented with 10ug/ml of blasticidin (Invitrogen, CA).

### Measurement of the body temperature by subcutaneous devices

Body temperature was measured using temperature-sensitive data loggers implanted in each animal of the needle infected groups, as previously described [35]. Experimental protocols were approved by the Animal Research Ethics Committee of the University of the Witwatersrand: 2015/07/31/C. Briefly, animals under sedation were injected with a local anaesthetic in the flank, the area was shaved, disinfected with chlorhexidine gluconate (Hibitane, SA) followed by an incision of the skin of about 5 cm. Data loggers were implanted below the skin and panniculus muscle into the flank and secured to the muscle with nylon sutures (NY924, size 0, SA). The surgical site was sutured closed with dissolvable sutures (Viamac VM514, size 2) and the surgery wounds sprayed with an antiseptic spray (Necrospray, Centaur Labs).

Figure 1 shows the variation of residuals through-out the experiment for SAT 1,2, 3 and the raw temperature for each individual. Adjusted body temperatures from the implanted animals before being exposed to FMDV were used to create a reference range (Additional figure 1). We fitted a nonlinear curve to the animals data over time so we could better include time in our assessment of body temperature. One animal had repeated outlier readings (as measured in a ROUT Analysis ^1^, Q=10%) so was omitted, resulting in n=11. A nonlinear curve with a sine function was fit to the data using robust nonlinear regression with constraints of amplitude >0, wavelength =1, frequency = 1 day and phase shift between 0 and 6.3 (2pi). The residuals were evaluated and found to vary between −1.057 and 1.042 (~39°C). The best fit values for this line are amplitude =0.492, wavelength =1 (constrained), Phaseshift = 3.378, frequency =1 and baseline temperature =38.29C. We calculated the residuals for the experimentally infected animals using the fitted nonlinear curve shown in Figure 1a with any value above a residual of 1.042 being considered a fever (above ~39C), and omitting any time point within 1 hour of a capture period. Using these residuals, we were able to calculate the length of each fever and the time it began. We also reported the peak temperature reached and the time point at which it was reached. To calculate the initial timepoint an animal mounted a fever and remove small fluctuations that may not be fever we took the time point at which at animal mounted a fever (first residual above 1.042) if the fever was sustained (all residuals above 1.042) for at least 6 consecutive hours. For the return to normal body temperature, we applied the same requirement, residuals had to be below 1.042 for at least 6 consecutive hours.

**Figure 1.**
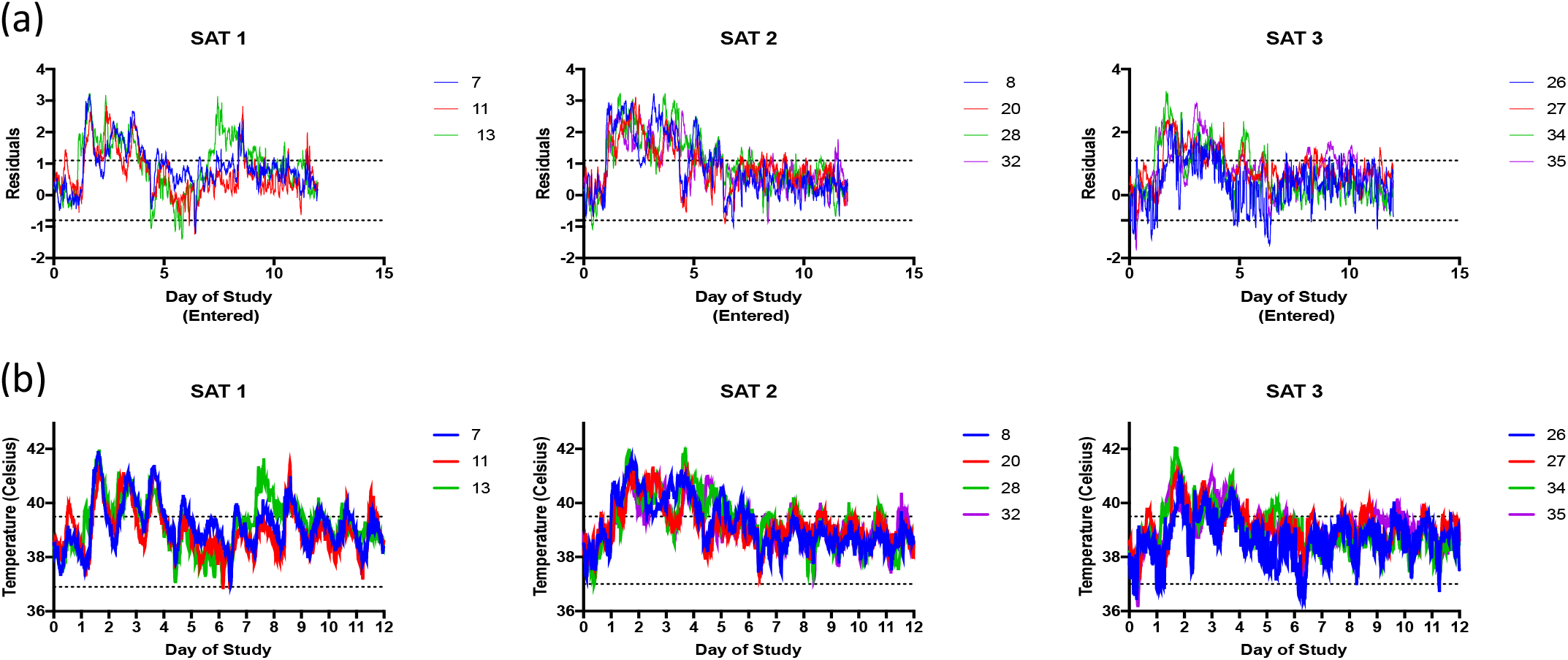
Body temperature in FMDV infected African buffalo. Body [7]temperature, measured by temperature-sensitive data loggers, of each individual buffalo after needle infection with FMDV SAT1, SAT2 or SAT3. Panel a represents the residuals from the fitted line, with a value above the dotted line considered a fever if the residuals stay above the line for 6 consecutive hours (72 readings). Panel b represents the raw temperatures over time. The dotted lines represent the normal temperatures as determined by the calculated reference range. Body temperature fluctuates during the day and all animals show high temperature at 1-1.7 dpi and remained elevated for between 2-5 dpi with a peak of maximum temperature of >41°C between 1.5-3.1 dpi. SAT2 infected animals have an elevated temperature and for longer duration compared to SAT1 and SAT3 (p 0.08 and p 0.016, respectively). All buffalo from the SAT1 group and 1 animal from SAT3 show a short second peak of high temperature at day 8 that lasted approximately 1 day only.

### FMDV RNA detection in serum, probang, nasal and tonsil swabs by reverse transcriptase qPCR (RT-qPCR)

RNA templates were extracted from 100ul of sample (serum, probang, nasal and tonsil swab) to a final elution volume of 80ul using MagNA Pure LC RNA isolation kit (Roche) and the KingsFisher Flex 96 robot (Thermofisher). Viral load was determined by means of RT-qPCR using primers targeting the conserved 3D^pol^-coding region of FMDV genome [36]. SAT serotype-specific primers and probes as previously described [10] were also used in tonsil swab samples. Forty cycles of PCR were carried out on a Stratagene Mx3005P QPCR system using MXPro MX3005 v3 software (Stratagene, UK). Cycle threshold (Ct) values were converted to FMDV genome copy number (GCN) by using a linear regression model with serial dilutions of *in vitro* synthetized RNA standard. Results were expressed as Log10 GCN/ml of sample. A cut-off of 1 GCN/5ul of RNA was used for all samples which resulted in detection thresholds of 2.2 log10 FMDV GCN/ml of sample.

### Air sampling

To investigate the possible aerosol transmission of FMDV in buffalo, we collected the aerosols exhaled by the NI buffalo using a Coriolis Air Sampler (Bertin Technologies) [37]. The Coriolis Air Sampler collects the aerosols in a plastic bottle filled with Eagles media with antibiotics that is connected to a high volume vacuum pump with an airflow rate of 300 liters/min. The air Sampler was positioned approximately 1 meter away from the mouth of the NI animals under sedation for 10 minutes, on days 0, 2, 4, 6 and 8 of the experiment. Aliquots of 1ml of media collected from the plastic bottle was analysed by RT-qPCR for the detection of viral particles.

### Virus isolation

Virus isolation from the oropharynx (OP) samples (probang and tonsil swab) and serum was performed in a monolayer of ZZR-127 goat epithelial cells following the procedures described by the Office International des Epizoties Manual of Diagnostic test (OIE Manual, 2021). When no cytopathic effect was observed after 48 hours of incubation a second passage of virus was performed on new ZZR-127. Positive cytopathic effects were confirmed for the presence of FMDV by RT-qPCR.

### Detection of FMDV neutralizing antibodies by virus neutralization test (VNT)

Serum samples were assayed for the presence of homologous neutralizing anti-FMDV antibodies by virus neutralization test (VNT) as described elsewhere_[38]. Briefly, 2-fold dilutions of serum are incubated with 100 TCID_50_ of SAT1, 2 or SAT3 FMDV in a monolayer of IB-RS-2 cells in 96well plates for three days. Number of wells with cytopathic effect (CPE) are counted and titres are expressed as the Log_10_ of the reciprocal of the highest dilution of serum that neutralized the virus in 50% of the wells. Titres >1.6 Log_10_ are considered to reach the threshold of protection according to Barnett and colleagues [39]

### Detection of FMDV antibodies against the non-structural proteins by ELISA

Serum samples were analysed for the detection of antibodies against the viral non-structural proteins (NSP). A PrioCHECK FMDV NS ELISA (Prionics^®^, The Netherlands) was performed according to the manufacturers’ specifications. Results more than 50% of percentage of inhibition (PI) are considered positive.

### Determination of type I/III IFN, TNF-α and IFN-ɣ in serum

An Mx/chloramphenicol acetyltransferase (Mx-CAT) reporter assay was used to determine the levels of biologically active IFN in serum samples (Fray et al., 2001). Briefly, serum samples were incubated on MDBK-t2 cells for 24h at 37°C and 5% CO2. Cells were then lysed in lysis buffer and CAT expression, induced by antiviral proteins present in the serum, was determined from the cell lysate using an ELISA kit (Roche) in accordance with the company instructions. Units of antiviral activity per ml of serum were calculated from a standard curve using recombinant bovine IFN-α [40]. A cut-off of 0.76 iu/ml was established by measuring the average of the basal levels plus 2 times the standard deviation.

The levels of TNF-α in buffalo sera were determined by means of ELISA using a commercial kit (RayBio ELB-TNF-α) according to the manufacture’s protocol. Results are expressed as μg /ml of serum. A cut-off of 1.76 μg/ml was established by measuring the average of the basal levels plus 2 times the standard deviation.

The quantitative determination of IFN-ɣ in buffalo serum was assayed by a commercial bovine IFN gamma sandwich ELISA test (Bio-Rad) following the manufacture’s specifications. Results are expressed as μg/ml extrapolated from a standard curve of recombinant bovine IFN-ɣ. A cut-off of 1.04 μg/ml was established by measuring the mean of the basal levels and adding 2 times the standard deviation of to the mean value.

### Determination of acute phase proteins in serum: serum amyloid A (SAA) and haptoglobin

Buffalo serum samples were tested for the levels of serum amyloid A (SAA) protein in a sandwich ELISA based on the instructions provided by the manufacturer (Life Diagnostics). Results are reported as ng/ml. A cut-off of 546 ng/ml was established by measuring the average of the basal levels plus 2 times the standard deviation for the single and co-infection experiments, respectively.

A commercial kit (Life Diagnostics, Inc) specific for bovine and based on a sandwich ELISA was used for the quantitative determination of haptoglobin in buffalo serum following the instructions. Results are expressed as ng/ml. A cut-off of 802 ng/ml was established by measuring the average of the basal levels plus 2 times the standard deviation.

### Statistical analysis

Data on maximum body temperature, peak, initial elevation, and duration of high temperature was analysed by R (version 3).

Virus load in serum and tonsil swab samples, FMDV immune response (VNT, TNF, Interferon ɣ and type I/III IFN and acute phase of proteins in serum (Haptoglobin and SAA) over time were analysed by determining, for each animal, the area under the curve (AUC), maximum value and day when the peak value was detected. For virus load (in serum and tonsil swab), VNT and NSP, first day with a positive value was also identified. Finally, duration of shedding was estimated for virus load in serum; duration was defined as the interval between the midpoint of first observation with a log_10_ value and the preceding negative observation and the midpoint of last observation with a log_10_ value and the subsequent negative observation). The response time was measured as time of the first positive value for each parameter minus the first day that FMDV is detected (presence of FMDV in tonsil, blood or nose).

All measurements were compared for NI animals and contact animals (regardless of the serotype) and different serotypes among needle infected animals and contact animals (corrected by time of exposure) using Kruskal Wallis test. Median, minimum and maximum values and the Kruskal-Wallis statistics of virus loads, serology and immunological values stratified by serotype (SAT1, SAT2 and SAT3) is shown in additional table 1. Median, minimum and maximum values and the Kruskal-Wallis statistics of virus loads, serology and immunological values stratified by method of infection (needle versus contact) is shown in additional table 2. Correlation between viremia levels and FMDV in tonsils was done by Spearman’s rank test.

## Results

### Transmission of FMDV from needle inoculated to in-contact buffalo

FMDV was transmitted readily from NI animals to all in-contact buffalo within the first week of being mixed. FMDV was first detected in serum and or tonsil and nasal swabs in all NI animals synchronous at 2 dpi; and as expected, FMDV detection in the in-contact buffalo was more variable within and between groups and delayed (p<0.014) compared to NI. Therefore, the analysis of the values of the immunological parameters in the in-contact groups accounted for the day that virus was first detected. The first detection of FMDV infection of the in-contact animals was not significantly different between groups challenged with the different serotypes (p=0.103). FMDV was not detected in any of the air samples collected adjacent to the NI animals after infection (data not shown).

Infection was delayed in one animal in the SAT3 in-contact group, with FMDV first detected on day 9, and was omitted from analysis due to the limited samples available post onset of infection.

### Clinical signs, body temperatures and leukocyte counts

After FMDV challenge there was no significant change in total white blood cell count (Additional figure 2) and only minor mouth lesions were seen in 3 out of 24 animals (two animals from SAT1 NI and one from SAT3 in-contact groups), at 6-11 dpi. Lesions consisted of small, rounded vesicles of around 4 mm diameter, in the upper dental pad. No lesions were observed in the coronary band or in the tongue, except for the needle tracks where the inoculation occurred.

As shown in figure 1 and table 1, body temperatures were elevated (>39.5°C) after infection (between 1-1.7 dpi, in all animals and remained elevated for between 2-5 dpi with a peak of maximum temperature of >41°C between 1.5-3.1 dpi. The SAT2 group showed a quicker response time to initial elevation (1 dpi) compared to 1.3 and 1.4 days for SAT1 and 3, respectively (p=0.008) and a longer duration of 5.3 days *versus* 3.01 and 2.96, for SAT1 and SAT3, respectively (p=0.016). All animals from SAT1 and one animal from SAT3 infected groups, showed a short second peak of pyrexia at 8 dpi that lasted approximately 1 day.

**Table 1.** Calculations of the residuals of body temperature for all African buffalo infected with SAT1, SAT2 and SAT3 over time; and its median values (minimum-maximum) and the Kruskal-Wallis statistics stratified by serotype.

| Group | Animal ID | Max Temp | Time of Peak (days) | Time of Initial Elevation (days) | Length of Fever in Days | Time of 2nd peak (days) | Length of Fever 2nd peak |
| --- | --- | --- | --- | --- | --- | --- | --- |
| SAT1 | 7 | 41.88 | 1.61 | 1.36 | 3.01 | 8.38 | 0.62 |
| SAT1 | 11 | 41.4 | 1.65 | 1.37 | 2.8 | 8.4 | 1.23 |
| SAT1 | 13 | 41.98 | 1.65 | 1.1 | 3.14 | 7 | 2.51 |
| Median values (minimum- maximum) |  | 41.88 (41.44-41.98) | 1.65 (1.61-1.65) | 1.36 (1.10-1.37) | 3.01 (2.8-3.14) |  |  |
| SAT2 | 8 | 41.04 | 3.18 | 1 | 5.32 | NA | NA |
| SAT2 | 20 | 41.34 | 2.53 | 1.09 | 3.27 | NA | NA |
| SAT2 | 28 | 42 | 1.66 | 1.01 | 5.31 | NA | NA |
| SAT2 | 32 | 41.26 | 1.57 | 1.01 | 5.32 | NA | NA |
| Median values (minimum- maximum) |  | 41.3 (41.04-42) | 2.09 (1.57-3.18) | 1.01 (1.0-1.09) | 5.32 (3.27-5.32) |  |  |
| SAT3 | 26 | 40.91 | 1.83 | 1.72 | 2.58 | NA | NA |
| SAT3 | 27 | 41.15 | 1.75 | 1.52 | 2.87 | NA | NA |
| SAT3 | 34 | 42.05 | 1.68 | 1.13 | 3.21 | 8.74 | 1.66 |
| SAT3 | 35 | 41.07 | 2.99 | 1.28 | 3.05 | NA | NA |
| Median values (minimum- maximum) |  | 41.11 (40.91-42.05) | 1.79 (1.68-2.99) | 1.4 (1.13-1.72) | 2.96 (2.5-3.21) |  |  |
|  | P-value | 0.47 | 0.164 | 0.0088 | 0.016 |  |  |

### Viral dynamics in blood, nasal swabs and oropharynx

Virus genome dynamics in serum samples from NI animals were comparable in all three groups (Figure 2a). The highest FMDV genome copy number was detected by 2 dpi in all the NI animals. Detection of virus genome in blood declined during 4-6 dpi and were undetectable by 8 dpi.

**Figure 2.**
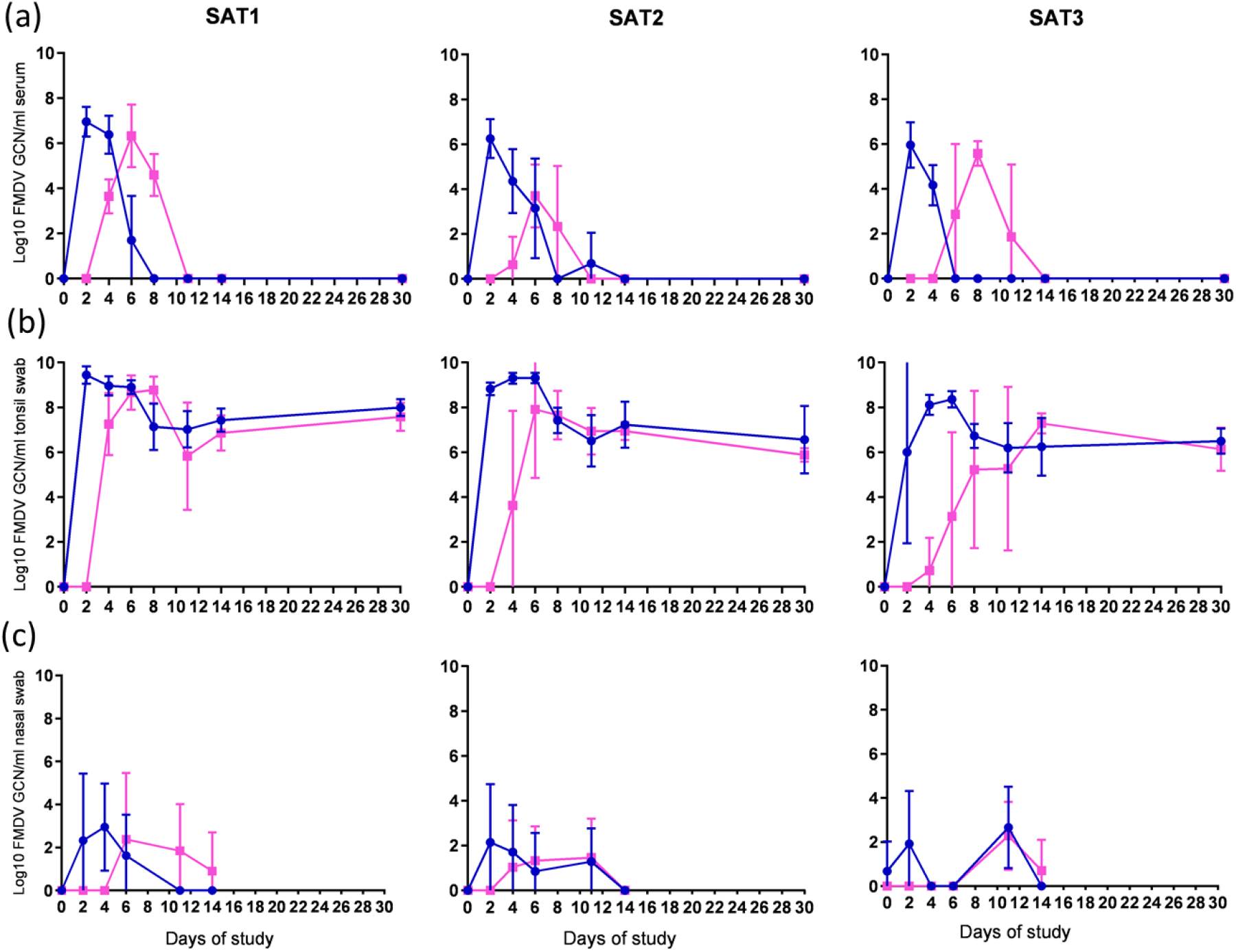
FMDV genome copy number dynamics in African buffalo. Detection of FMDV genome copy numbers (GCN) by qRT-PCR in serum (a), tonsil swab (b) and nasal swab (c) from animals needle infected with SAT1, SAT2 and SAT3 FMDV serotypes (blue lines, round symbol) *versus* contact challenge (pink, square symbol). NI animals were challenged on day 0 of the study, and contact animals were mixed with NI groups on day 2. Graphs represent the mean and SEM for each group at each time point.

Virus genome dynamics in serum from in-contact animals was more variable within each group and differences were observed between serotypes. The SAT2 in-contact group showed significantly lower genome copy numbers in serum with averages of 2.57 GCN/ml *versus* 3.24 and 3.42 GCN/ml for SAT3 and SAT1, respectively (p<0.028), while virus genome was detected earlier in the SAT1 in-contact animals compared to the SAT2 and SAT3 animals (p = 0.017). Also, the duration of detectable genome in serum was longer for the SAT1 group (6.5 days) compared to the SAT2 (4.5 days) and SAT3 (2.5 days) animals (p = 0.021).

FMDV genome could be detected in the oropharynx (OP) at 2 dpi regardless of the route of infection, (Figure 2b). In general, GCN values peak between 3 to 6 dpi, except for SAT3 in-contact animals which showed a significant delay (5 to 12dpi) (p<0.045). FMDV genome was detected in all tonsil swabs until day 30 of the experiment. NI animals had higher GCN in OP compared to in-contact infected animals from day 2 to day 30 of the experiment (p=0.004), with the SAT1 NI group showing higher values (5.43 GCN/ml), compared to SAT2 and SAT3 NI groups (5.39 GCN/ml and 5.27 GCN/ml) (p=0.048). By 30 dpi, tonsil swabs were analysed by qRT-PCR using SAT specific primers (Table 2) and results indicated that no evidence of cross-infection was detected in any of the groups housed separately during the experiment.

**Table 2.** Comparison of FMDV genome (Log_10_ GCN/ml) and virus isolation (indicated with an underline) in serum, probang and tonsil swab along the study measured by 3D qRT-PCR and by SAT specific qRT –PCR in tonsil swab on day 30. Cut-off was established at 1 GCN/5ul of RNA. Those samples that were FMDV isolation positive are indicated with an underline.

|  |  |  | Log<sub>10</sub> FMDV GCN/ml serum |  |  |  |  |  | Log<sub>10</sub> FMDV GCN/ml probang |  |  | Log<sub>10</sub> FMDV GCN/ml tonsil swab |  |  | Log<sub>10</sub> FMDV SAT specific GCN/ml tonsil swab at day 30 |  |  |
| --- | --- | --- | --- | --- | --- | --- | --- | --- | --- | --- | --- | --- | --- | --- | --- | --- | --- |
| FMDV challenge | Group (FMDV strain) | ID animal | Day 2 | Day 4 | Day 6 | Day 8 | Day 11 | Day 14 | Day 8 | Day 14 | Day 30 | Day 8 | Day 14 | Day 30 | SAT1 | SAT2 | SAT3 |
| Needle infected (NI) | SAT1 | 7 | 7.67 | <u>7.36</u> | 3.53 | 0.00 | 0.00 | 0.00 | <u>5.88</u> | <u>5.39</u> | 5.01 | <u>7.62</u> | <u>7.28</u> | <u>8.07</u> | 8.51 | 0.00 | 0.00 |
|  | SAT1 | 10 | <u>7.35</u> | 5.56 | 0.00 | 0.00 | 0.00 | 0.00 | <u>5.13</u> | <u>5.89</u> | 3.99 | <u>7.64</u> | <u>7.10</u> | <u>8.05</u> | 8.25 | 0.00 | 0.00 |
|  | SAT1 | 11 | <u>6.31</u> | 5.80 | 0.00 | 0.00 | 0.00 | 0.00 | <u>4.91</u> | <u>4.27</u> | <u>4.56</u> | <u>7.71</u> | <u>8.21</u> | <u>8.38</u> | 8.90 | 0.00 | 0.00 |
|  | SAT1 | 13 | <u>6.49</u> | <u>6.79</u> | 3.27 | 0.00 | 0.00 | 0.00 | <u>3.93</u> | <u>5.08</u> | <u>3.95</u> | <u>5.59</u> | <u>7.14</u> | <u>7.47</u> | 7.92 | 0.00 | 0.00 |
|  | SAT2 | 8 | <u>6.40</u> | 3.84 | 0.00 | 0.00 | 0.00 | 0.00 | <u>5.04</u> | 0.00 | 0.00 | <u>7.28</u> | <u>7.63</u> | <u>7.68</u> | 0.00 | 7.21 | 0.00 |
|  | SAT2 | 20 | <u>7.31</u> | 5.91 | 3.50 | 0.00 | 0.00 | 0.00 | <u>4.53</u> | <u>4.40</u> | 0.00 | <u>8.03</u> | <u>8.43</u> | 5.76 | 0.00 | 5.67 | 0.00 |
|  | SAT2 | 28 | <u>5.20</u> | <u>2.63</u> | 3.85 | 0.00 | 2.75 | 0.00 | <u>5.97</u> | <u>2.94</u> | 3.82 | <u>6.70</u> | <u>6.80</u> | 4.85 | 0.00 | 5.32 | 0.00 |
|  | SAT2 | 32 | 6.09 | 5.05 | 5.22 | 0.00 | 0.00 | 0.00 | <u>6.42</u> | 3.81 | 0.00 | <u>7.67</u> | 6.04 | <u>7.96</u> | 0.00 | 7.95 | 0.00 |
|  | SAT3 | 26 | <u>7.46</u> | 3.20 | 0.00 | 0.00 | 0.00 | 0.00 | <u>5.74</u> | <u>5.03</u> | 3.32 | <u>7.31</u> | <u>7.44</u> | <u>7.27</u> | 0.00 | 0.00 | 8.04 |
|  | SAT3 | 27 | <u>5.46</u> | <u>5.38</u> | 0.00 | 0.00 | 0.00 | 0.00 | 3.42 | 2.39 | 2.97 | <u>6.44</u> | 4.42 | 6.00 | 0.00 | 0.00 | 6.85 |
|  | SAT3 | 34 | 5.22 | 4.09 | 0.00 | 0.00 | 0.00 | 0.00 | <u>4.64</u> | <u>4.04</u> | 3.51 | <u>7.04</u> | <u>6.52</u> | <u>6.51</u> | 0.00 | 0.00 | 6.87 |
|  | SAT3 | 35 | <u>5.70</u> | 4.01 | 0.00 | 0.00 | 0.00 | 0.00 | <u>3.63</u> | 3.12 | 3.35 | <u>6.14</u> | <u>6.59</u> | <u>6.19</u> | 0.00 | 0.00 | 8.40 |
| In-contact | SAT1 | 2 |  | 3.32 | <u>5.87</u> | <u>5.79</u> | 0.00 | 0.00 | <u>4.59</u> | <u>3.20</u> | 0.00 | <u>6.31</u> |  |  |  |  |  |
|  | SAT1 | 4 |  | 2.74 | 5.55 | 4.84 | 0.00 | 0.00 | <u>5.66</u> | <u>5.29</u> | <u>4.89</u> | <u>8.93</u> | <u>6.87</u> | <u>7.29</u> | 7.82 | 0.00 | 0.00 |
|  | SAT1 | 19 |  | 4.41 | 5.51 | 4.03 | 0.00 | 0.00 | <u>6.34</u> | <u>4.30</u> | 0.00 | <u>8.96</u> | <u>7.62</u> | <u>7.15</u> | 8.06 | 0.00 | 0.00 |
|  | SAT1 | 33 |  | 4.10 | <u>8.39</u> | 3.72 | 0.00 | 0.00 | <u>5.33</u> | <u>5.27</u> | <u>5.15</u> | <u>9.30</u> | <u>7.18</u> | <u>8.30</u> | 8.65 | 0.00 | 0.00 |
|  | SAT2 | 5 |  | 2.50 | 2.38 | 0.00 | 0.00 | 0.00 | <u>4.44</u> | <u>3.69</u> | 0.00 | <u>8.89</u> | <u>6.53</u> | 5.86 | 0.00 | 5.33 | 0.00 |
|  | SAT2 | 9 |  | 0.00 | <u>5.64</u> | 4.80 | 0.00 | 0.00 | <u>4.36</u> | 4.24 | 0.00 | <u>8.24</u> | <u>7.49</u> | 6.25 | 0.00 | 6.15 | 0.00 |
|  | SAT2 | 22 |  | 0.00 | 3.08 | 4.55 | 0.00 | 0.00 | <u>4.57</u> | 0.00 | 0.00 | <u>6.84</u> | <u>6.83</u> | 5.89 |  |  |  |
|  | SAT2 | 29 |  | 0.00 | 3.71 | 0.00 | 0.00 | 0.00 | <u>5.22</u> | 4.53 | 3.33 | <u>6.66</u> | <u>6.92</u> | <u>5.50</u> | 0.00 | 5.41 | 0.00 |
|  | SAT3 | 12 |  | 0.00 | 0.00 | <u>5.37</u> | 5.59 | 0.00 | <u>4.64</u> | <u>4.44</u> | <u>4.20</u> | <u>7.23</u> | <u>7.60</u> | <u>5.68</u> | 0.00 | 0.00 | 6.25 |
|  | SAT3 | 15 |  | 0.00 | 0.00 | 0.00 | 0.00 | 5.42 | 0.00 | <u>3.96</u> | <u>4.88</u> |  | <u>6.74</u> | 5.63 | 0.00 | 0.00 | 6.59 |
|  | SAT3 | 16 |  | 0.00 | 2.38 | <u>6.20</u> | 0.00 | 0.00 | <u>4.93</u> | 2.87 | <u>3.92</u> | <u>7.26</u> | <u>7.12</u> | <u>7.57</u> | 0.00 | 0.00 | 8.40 |
|  | SAT3 | 17 |  | 0.00 | <u>6.22</u> | 5.19 | 0.00 | 0.00 | 0.00 | 3.53 | 4.30 | <u>6.42</u> | <u>7.70</u> | 5.67 | 0.00 | 0.00 | 6.65 |
| # of samples RT qPCR positive |  |  | 12 | 17.00 | 15.00 | 9.00 | 2.00 | 1.00 | 23.00 | 22.00 | 16.00 | 23.00 | 23.00 | 23.00 |  |  |  |
| # of samples RT qPCR and VI positive |  |  | 8 | 3.00 | 3.00 | 3.00 |  | 0.00 | 22.00 | 15.00 | 8.00 | 23.00 | 22.00 | 15.00 |  |  |  |
| VI positive (%) |  |  | 66.7 | 17.6 | 20.0 | 33.3 | 0.0 | 0.0 | 95.7 | 68.2 | 50.0 | 100.0 | 95.7 | 65.2 |  |  |  |
| Averages VI (%) |  |  |  |  | 22.9 |  |  |  |  | 71.3 |  |  | 87 |  |  |  |  |

Virus genome was first detected in nasal swabs on 2 and 6.5 dpi (group mean values) in NI and in-contact groups, respectively (Figure 2c). Most of the animals were negative by 14 dpi. Contrary to the high GCN in blood and OP, virus genome detection in nasal swabs was intermittent and reached maximum values of 3.4 and 3.34 GCN/ml in NI and in-contact groups, respectively. No statistical differences in the dynamics of shedding in nasal swabs were observed between groups.

Serum, probang, and tonsil and nasal swabs were also analysed by virus isolation (VI) from 2-30 dpi of the experiment (Table 2). On day 2 and 4 after virus exposure, only 66% and 17% respectively of the serum samples were positive for virus isolation which contrasts with the high qRT-PCR values in all serum samples on these days. Virus was isolated from 87% and 71% of the qRT-PCR positive samples from tonsil swab and probang, respectively, on days 8, 14 and 30 of the experiment. Also, the mean GCN from all VI positive samples was higher in tonsil swabs (p<0.001) (Additional figure 3) thus indicating that tonsil swab is the most reliable method for detecting FMD live virus and genome in African buffalo. On day 30 of the experiment infectious virus was isolated from tonsil swabs and/or probang from 16 (9 NI and 7 in contact), out of 24 (66.6%) infected animals (Table 3), however there was no association of level of viral loads in oropharynx or route of virus exposure with the carrier status (p=0.33 and P=0.553, respectively). Infectious virus could not be isolated from any nasal secretions.

**Table 3.** Number of animals with vesicles and number of animals that were carrier by day 30 of the study

|  | NI |  |  |  | Contact |  |  |  | p-value |
| --- | --- | --- | --- | --- | --- | --- | --- | --- | --- |
|  | SAT1 | SAT2 | SAT3 | Total NI (%) | SAT1 | SAT2 | SAT3 | Total contact (%) |  |
| Clinical signs (vesicles) | 2 | 0 | 0 | 2 (16.7) | 0 | 0 | 1 | 1 (8.3) | 1 |
| Carrier at day 30 (VI+) | 4 | 2 | 3 | 9 (75) | 3 | 1 | 3 | 7 (58.3) | 0.556 |

### Humoral immune response to FMDV

The specific humoral immune responses induced by the different FMDV SAT serotypes after NI were not significantly different, however differences are observed in in contact groups. (Figure 3a). In general, FMDV infected buffalo developed virus neutralizing antibody titres (VNTs) within 2 to 6 days post virus exposure. VNTs rapidly increased after first detection and were maintained at their maximum titres until the end of the study on day 30. The route of infection did not influence the magnitude of the VNTs but NI reached protective titres faster compared to contact (p<0.002) and among the contact animals, the onset of the response was faster in SAT1 group (p<0.021), showing comparable levels with NI animals.

**Figure 3.**
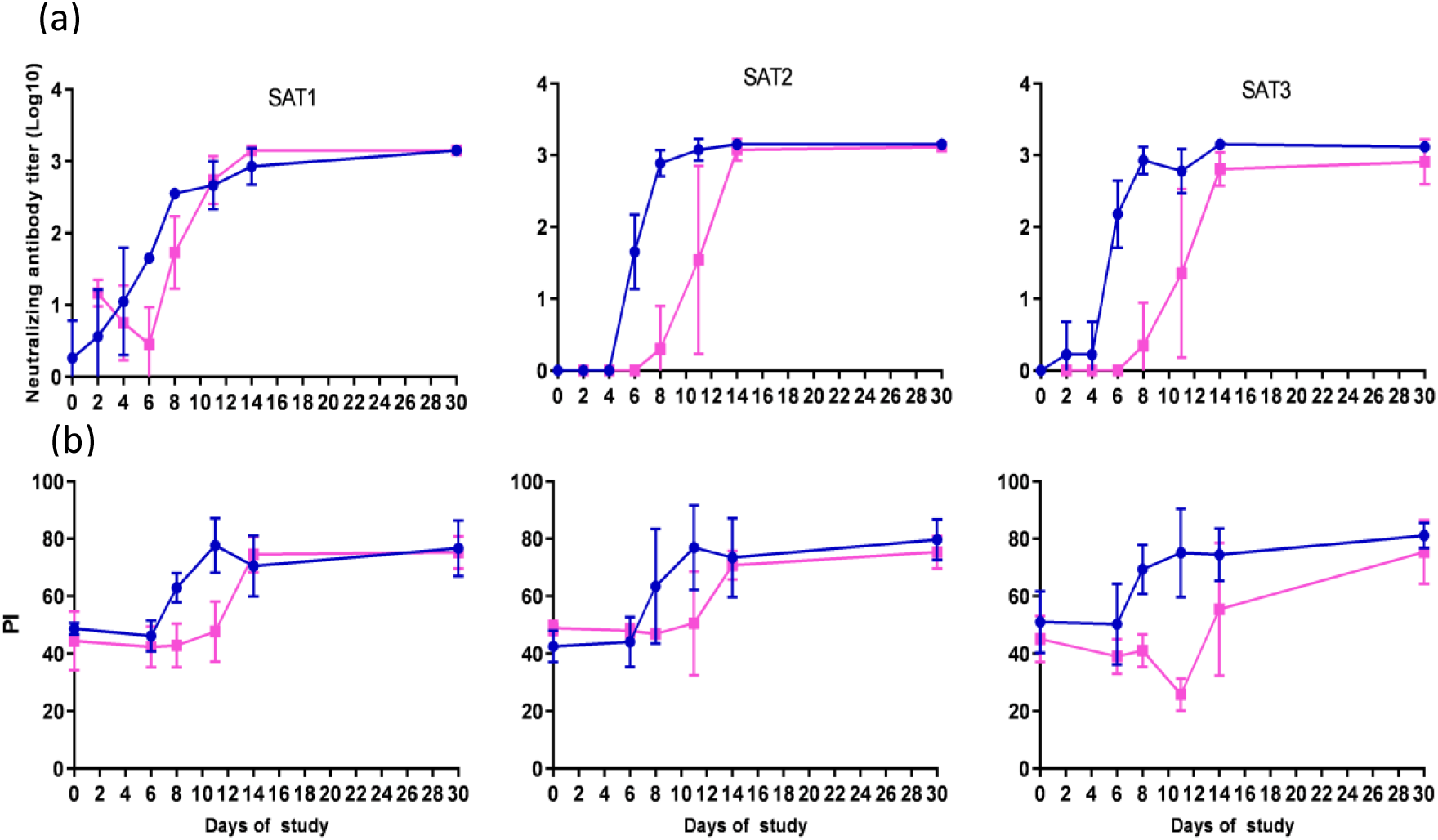
Specific humoral immune response induced by FMDV infection. Neutralizing antibody titers in log10 (a) and presence of antibody levels against FMDV non-structural proteins (NSP) (b), induced by SAT1, SAT2 and SAT3 FMDV infected animals by needle infection (blue lines, round symbol) or by contact challenge (pink, square symbol). NSP results are expressed by percentage of inhibition (PI) and cut-off is determined at 50% (dash line). Graphs represent the mean and SEM for each group at each time point.

Antibodies against the non-structural proteins (NSP) of FMDV were first detected at 8 dpi for the NI groups and significantly delayed in the in-contact groups (12 dpi, p = 0.001). The NSP antibody titres remained consistently elevated until 30 dpi (Figure 3b).

### Levels of Haptoglobin and SAA in serum of FMDV infected buffalo

Serum amyloid A (SAA) and haptoglobin were detected during acute FMD infection in buffalo. High concentrations of SAA were detected in serum of all animals immediately after virus infection (Figure 4a). Serum concentrations rapidly increased and peaked by 4-6 dpi. Levels declined progressively after the peak and by 14 dpi SAA levels were undetectable. While the total SAA response was not different across serotypes and route of infection, the induction of SAAs was delayed in in-contact animals (p<0.004), however, their peak levels were higher compared to NI (p<0007).

**Figure 4.**
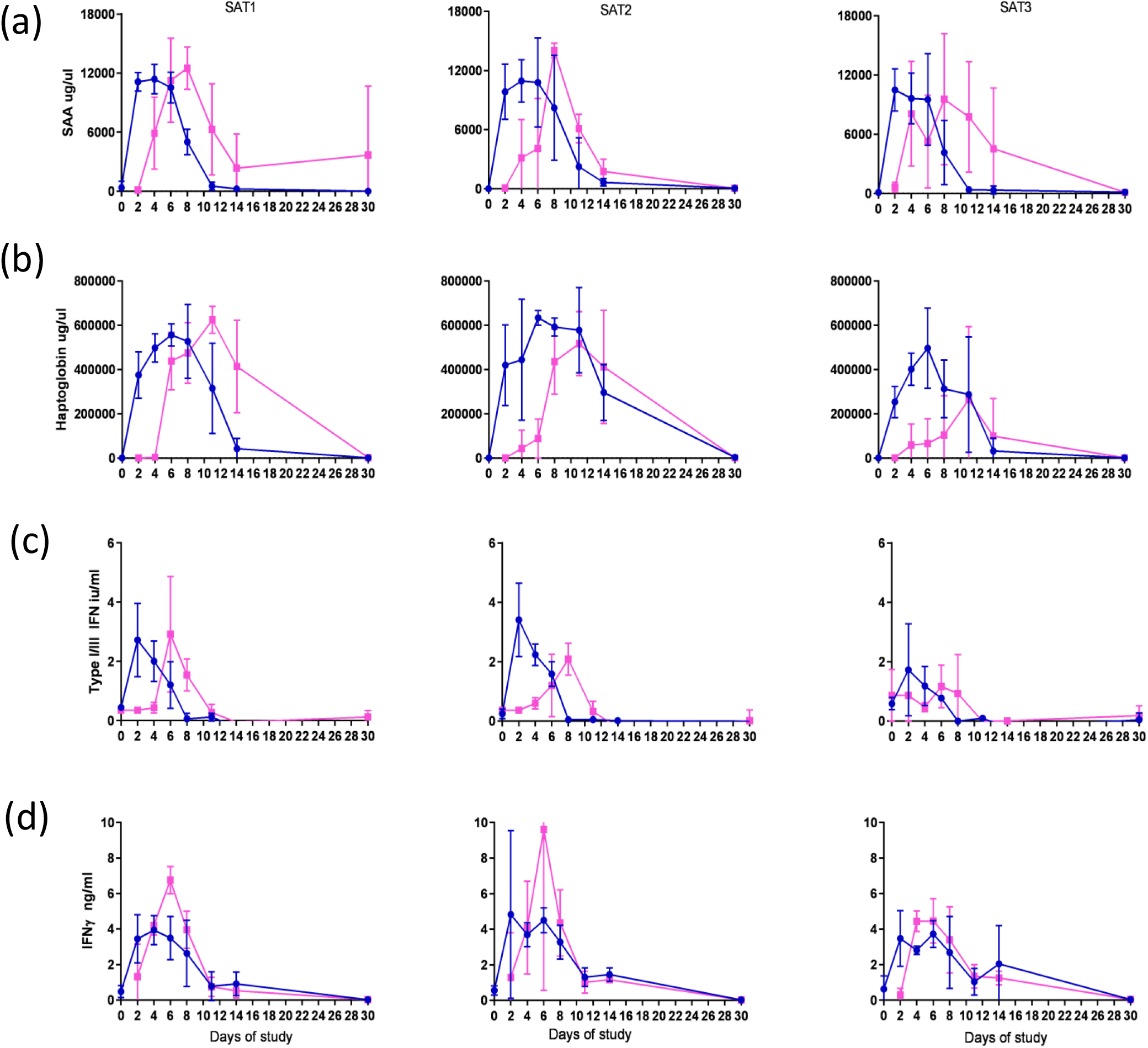
Innate immune response and acute phase proteins induced by FMDV in African buffalo. Concentrations of SAA (a), hatoglobin (b), type I/III IFN (c) and IFNɣ (d) in serum from animals infected with SAT1, SAT2 or SAT3 FMDV serotypes by NI (blue lines, round symbol) or contact challenge (pink lines, square symbol). Graphs represent the mean and SEM for each group at each time point. Cut-offs for SAA and hatoglobin were established at 546 and 802 ng/ul and cut off for type I IFN and IFNɣ was established at 0.76 iu/ml and 1.04 ng/ml, respectively.

Similar dynamics were observed in the concentration of haptoglobin in serum (Figure 4b) although levels were maintained for longer than SAA; by 30 dpi haptoglobin levels in all animals were normal. No differences in haptoglobin in serum were observed after the different routes of infection, but the magnitude of the response was the highest within the SAT2 NI groups (p<0.018).

### Innate immune response induced in FMDV infected buffalo

The dynamics of Type I/III IFNs and IFNɣ in serum were very similar in response to all SAT infections (Figure 4c). TypeI/III IFN were detected in the serum for approximately 6 dpi with a peak of 2 dpi for NI and 6 days after in-contact challenge (p=0.013). Similarly, the induction of IFNɣ was detected at 2 dpi and peaked at 6 dpi with maximum values for in-contact animals higher than NI (p=0.036) (Figure 4d). No differences in dynamics of TypeI/III IFN and IFNγ antiviral cytokines were found across serotypes, however some animals from the in-contact group had detectable levels to these cytokines even before FMDV was detected (p=0.031).

TNFα could not be consistently detected in FMDV infected buffalo (data not shown).

## Discussion

This study represents the most complete characterization of viral dynamics and immune responses to FMDV infection in African buffalo. FMD in African buffalo is generally regarded as mild or asymptomatic, since no (or very few) vesicles are observed even after a high dose of FMDV challenge [10, 41, 42]. Consistent with these previous reports, small vesicles restricted to the dental pad were only observed in two SAT1 NI buffaloes and one SAT3 in-contact challenged buffalo, contrary to cattle that present with vesicles at multiple sites, generally on the feet and tongue, after the onset of fever [1]. However, using temperature loggers, we demonstrated for the first time that African buffalo are indeed systemically affected by FMDV and develop consistent pyrexic responses early after needle infection (1-2 dpi) that last for approximately 3 to 5 days. Recent results highlighted that cattle with FMDV are substantially less likely to be infectious before showing clinical signs, including pyrexia and a significant increase of greater than 1°C in body temperature has been considered to be a good indicator of the onset of FMD clinical signs after experimental challenge [8] In fact, temperature has also been considered a good correlate of transmission of FMDV [9]]. In the absence of FMD lesions body temperature could probably be the most important correlate of transmission in African buffalo. Interestingly, SAT2 NI showed an earlier increased temperature and for longer compared to SAT1 and SAT3 infected animals, however the increased temperature was not associated with a higher virus load in serum or virus replication in the oropharynx. Within one week of FMDV exposure, buffalo also showed high levels of SAA in serum, similar to the profile detected in cattle [18]. Interestingly, SAT2 NI animals showed higher levels of haptoglobin in serum (p<0.018) compared to SAT1 and SAT3 challenged animals. Acute phase proteins are non-specific markers of inflammation, and although most buffalo did not show FMD lesions, they were all systemically affected by virus infection. Therefore, these proteins could be used as a surrogate marker of FMDV infection in African buffalo, as previously suggested [19].

It has been described that FMDV in cattle is highly contagious and R_0_ have been estimated to be between 21-88, [4, 6] even though the infectious period is brief (1.7 (0.3-4.8) days) [9]. Moreover, in domestic cattle there is a positive association between transmission, and presence of virus in air, and the onset of FMD clinical signs [8]. In this study, despite the lack of vesicles, and the absence of virus in air samples, all contact buffalo were readily infected after being in contact with the NI animals. Indeed, high levels of virus and virus genome were detected in the palatine tonsils by qRT-PCR and virus was isolated during the first 4-6 days after infection in all NI animals. These results indicate that the tonsils might be the main source of infectious virus in buffalo rather than vesicular lesions as described for cattle [9]. When comparing both types of pharyngeal samples, higher viral genome copies were detected by qRT-PCR in tonsil swabs compared to probang (p<0.001); these results corroborate previous findings suggesting that tonsil swabs performed better than probang for FMDV diagnosis [10]. Viral loads in tonsil decreased over time; however, most of the animals still were shedding virus by day 30 of the experiment, therefore, with potential of still transmitting FMDV by the end of the experiment.

FMDV was first detected concomitantly in tonsil swabs and blood from most of the animals within the first week after FMDV infection, only three animals from the in-contact group showed earlier detection in tonsil swabs than blood, in contrast, one animal from the NI group showed FMDV in blood before tonsils. It has been reported in cattle that virus detected in oropharynx provides the earliest indication of infection; but virus in the blood and nasal fluid may also be good candidates for preclinical indicators of infectiousness when virus levels exceed certain thresholds [8]. In this study, the presence of virus genome in nasal swabs was not easily detected and not consistent within groups (5 out of 24 animals were negative at all time points).

Virus genome could be detected in blood from infected buffalo for approximately 4 to 6 dpi, which is a longer duration than measured in cattle (2-4 dpi) [1, 9, 43]. Viral genome in blood correlated closely with the detection of viral genome in tonsil swabs until the appearance of neutralizing antibodies. Soon after neutralizing antibodies were detected the virus was cleared completely from the bloodstream, around 6dpi. Similar to cattle, FMDV detection in the oropharynx or tonsil is not affected by the presence of neutralizing antibodies [12]. High FMDV genome copy numbers were maintained in palatine tonsil until late after infection when the titres of neutralizing antibodies were maximum. In fact, by 30 dpi, FMDV could be isolated from tonsil swabs and/or probang in 16 out of 24 animals and these were identified as carriers. We and others have demonstrated that buffalo can remain persistently infected with FMDV for months and years ( [10, 24, 44]) and although transmission from carrier buffalo to naïve is difficult to reproduce ([41, 45] a recent publication demonstrated that it is indeed the inclusion of occasional transmission from carriers that rescues FMDV from extinction in isolated African buffalo populations ([7] In our study, the development of carriers did not correlate with clinical signs or acute host responses, as suggested for cattle [18]. There was also no association between the carrier status and infection route or serotype.

Altogether, these results demonstrated similar dynamics of FMDV infection and immune responses after needle infection or direct in-contact challenge in African buffaloes compared to cattle, despite marked differences in the clinical outcome [10, 18]. The reasons for the different clinical outcomes between the host species remains unclear, in addition to the lack of understanding of the mechanisms responsible for the tissue distribution of FMD vesicles in cattle [46],. We have demonstrated for the first time that pyrexia is a consistent clinical sign of FMD in African buffalo. In general, needle challenge leads to a synchronous, faster and higher viral loads in blood and oropharynx and specific humoral immune response while the innate and acute immune responses were similar in needle and in-contact challenged buffaloes. These differences could be explained by the variable time and the lower dose of infection in the in-contact group compared to high doses of virus in NI. The SAT1 virus was detected more rapidly after challenge compared to the SAT2 and SAT3 viruses and transmitted more readily to naïve buffalo. These results agree with our previous studies where we showed during mixed infections in individual buffalo, over time SAT1 persisted for longer periods compared to SAT2 and SAT3 viruses. [10, 47, 48]. The results are also consistent with our observation during a long-term study of an isolated buffalo herd that demonstrated SAT1 viruses persist more readily in a population [7]

These data provide important information to help understand the marked clinical differences between cattle and African buffalo in their response to FMDV infection. We have demonstrated that the typically mild clinical signs in African buffalo are not because virus replication or shedding are controlled and are not associated with a suppressed immune response to FMDV. We have also demonstrated that naïve buffaloes kept in contact with acutely infected buffaloes are readily infected despite the absence of high titre virus in vesicular fluids or lesions. Further studies are required to investigate cell mediated immune responses, and to determine if this arm of the immune response is accountable for the markedly different clinical outcomes in African buffalo compared to cattle. These data form a foundation for modelling the interplay of viral and immune response dynamics within African buffalo host and understanding the pathogenesis of these highly contagious viruses in populations of their natural reservoir host.

## Abbreviations

FMD: Foot and mouth disease
FMDV: foot and mouth disease virus
SAT: Southern Africa Territories
SAA: serum amyloid A
APP: acute phase proteins
OP: oropharynx
VNT: virus neutralizing test
MAbs: monoclonal antibodies
KNP: Kruger National Park
dpi: days post infection
NSP: non-structural protein
NI: needle infected
Co: contact infected
GCN: genome copy number
PI: percentage of inhibition
IFN: interferon
TNF: tumor necrosis factor

## Acknowledgements

Authors thank Dave Cooper and the boards of the South African national parks for supplying the FMDV free buffalo used in these studies, and the animal unit staff at Kruger National Park in South Africa for their invaluable assistance with the *in vivo* experiment. We also thank J. Gonzalez for helping to plan the experiment, B. Martinez for believing in our purpose and L. Stevenson from the animal facilities at The Pirbright Institute for her excellent assistance with the inventory of the samples.

## Author’s contributions

EPM, BB, BC, AJ, NJ and FM conceived and planned the experiments. EPM, BB, FZ, BD, FM, LKL, KS, AJ, AH, LS and BC carried out the experiments. EPM, BB, FZ and KS generated the data. EPM, GLV, BB and AJ analysed the data and contributed to the interpretation of the results. EPM and BC took the lead in writing the manuscript. All authors provided critical feedback and reviewed the manuscript.

## Availability of data and materials

The datasets analyzed during the current study are available from the corresponding authors upon request.

## Ethics approval

Experimental protocols were approved by the Animal Ethical Committee of the Department of Agriculture, Land Reform and Rural Development (DALRRD), KNP-BC-02 and SANParks N013-12.

## Authors details

^1^ The Pirbright Institute, Woking, Surrey, United Kingdom;^2^ Oregon State University, Corvallis, Portland, USA; ^3^ Agricultural Research Council of South Africa, Onderstepoort Veterinary Institute-Transboundary Animal Disease section (OVI-TAD), Vaccine and Diagnostic Development Programme, Onderstepoort, Gauteng, South Africa.; ^4^ State Veterinary Services, P.O. Box 12, Skukuza, 1350; ^5^ UCL Institute of Prion Diseases, London, United Kingdom; ^6^ Brain Function Research Group, School of Physiology, Faculty of Health Sciences, University of the Witwatersrand, Johannesburg, South Africa; ^7^ South Africa Bill and Melinda Gates Foundation, Seatle, USA

## References

1. Grubman MJ, Baxt B: Foot-and-mouth disease. Clin Microbiol Rev 2004, 17(2):465–493.

2. Casey-Bryars M, Reeve R, Bastola U, Knowles NJ, Auty H, Bachanek-Bankowska K, Fowler VL, Fyumagwa R, Kazwala R, Kibona T et al: Waves of endemic foot-and-mouth disease in eastern Africa suggest feasibility of proactive vaccination approaches. Nature ecology & evolution 2018, 2(9):1449–1457.

3. Brito BP, Rodriguez LL, Hammond JM, Pinto J, Perez AM: Review of the Global Distribution of Foot-and-Mouth Disease Virus from 2007 to 2014. Transboundary and emerging diseases 2017, 64(2):316–332.

4. Chis Ster I, Dodd PJ, Ferguson NM: Within-farm transmission dynamics of foot and mouth disease as revealed by the 2001 epidemic in Great Britain. Epidemics 2012, 4(3):158–169.

5. Eblé PL, Orsel K, van Hemert-Kluitenberg F, Dekker A: Transmission characteristics and optimal diagnostic samples to detect an FMDV infection in vaccinated and non-vaccinated sheep. Veterinary microbiology 2015, 177(1-2):69–77.

6. Hayer SS, VanderWaal K, Ranjan R, Biswal JK, Subramaniam S, Mohapatra JK, Sharma GK, Rout M, Dash BB, Das B et al: Foot-and-mouth disease virus transmission dynamics and persistence in a herd of vaccinated dairy cattle in India. Transboundary and emerging diseases 2018, 65(2):e404–e415.

7. Jolles A, Gorsich E, Gubbins S, Beechler B, Buss P, Juleff N, de Klerk-Lorist LM, Maree F, Perez-Martin E, van Schalkwyk OL et al: Endemic persistence of a highly contagious pathogen: Foot-and-mouth disease in its wildlife host. Science (New York, NY) 2021, 374(6563):104–109.

8. Chase-Topping ME, Handel I, Bankowski BM, Juleff ND, Gibson D, Cox SJ, Windsor MA, Reid E, Doel C, Howey R et al: Understanding foot-and-mouth disease virus transmission biology: identification of the indicators of infectiousness. Veterinary research 2013, 44:46.

9. Charleston B, Bankowski BM, Gubbins S, Chase-Topping ME, Schley D, Howey R, Barnett PV, Gibson D, Juleff ND, Woolhouse ME: Relationship between clinical signs and transmission of an infectious disease and the implications for control. Science (New York, NY) 2011, 332(6030):726–729.

10. Maree F, de Klerk-Lorist LM, Gubbins S, Zhang F, Seago J, Perez-Martin E, Reid L, Scott K, van Schalkwyk L, Bengis R et al: Differential Persistence of Foot-and-Mouth Disease Virus in African Buffalo Is Related to Virus Virulence. Journal of virology 2016, 90(10):5132–5140.

11. Arzt J, Juleff N, Zhang Z, Rodriguez LL: The pathogenesis of foot-and-mouth disease I: viral pathways in cattle. Transboundary and emerging diseases 2011, 58(4):291–304.

12. Stenfeldt C, Eschbaumer M, Rekant SI, Pacheco JM, Smoliga GR, Hartwig EJ, Rodriguez LL, Arzt J: The Foot-and-Mouth Disease Carrier State Divergence in Cattle. Journal of virology 2016, 90(14):6344–6364.

13. Perez-Martin E, Weiss M, Diaz-San Segundo F, Pacheco JM, Arzt J, Grubman MJ, de los Santos T: Bovine type III interferon significantly delays and reduces the severity of foot-and-mouth disease in cattle. Journal of virology 2012, 86(8):4477–4487.

14. Perez-Martin E, Diaz-San Segundo F, Weiss M, Sturza DF, Dias CC, Ramirez-Medina E, Grubman MJ, de los Santos T: Type III interferon protects swine against foot-and-mouth disease. Journal of interferon & cytokine research : the official journal of the International Society for Interferon and Cytokine Research 2014, 34(10):810–821.

15. Dias CC, Moraes MP, Weiss M, Diaz-San Segundo F, Perez-Martin E, Salazar AM, de los Santos T, Grubman MJ: Novel antiviral therapeutics to control foot-and-mouth disease. Journal of interferon & cytokine research : the official journal of the International Society for Interferon and Cytokine Research 2012, 32(10):462–473.

16. Medina GN, Segundo FD, Stenfeldt C, Arzt J, de Los Santos T: The Different Tactics of Foot-and-Mouth Disease Virus to Evade Innate Immunity. Front Microbiol 2018, 9:2644.

17. Moraes MP, de Los Santos T, Koster M, Turecek T, Wang H, Andreyev VG, Grubman MJ: Enhanced antiviral activity against foot-and-mouth disease virus by a combination of type I and II porcine interferons. Journal of virology 2007, 81(13):7124–7135.

18. Stenfeldt C, Heegaard PM, Stockmarr A, Tjørnehøj K, Belsham GJ: Analysis of the acute phase responses of serum amyloid a, haptoglobin and type 1 interferon in cattle experimentally infected with foot-and-mouth disease virus serotype O. Veterinary research 2011, 42(1):66.

19. Glidden CK, Beechler B, Buss PE, Charleston B, de Klerk-Lorist LM, Maree FF, Muller T, Pérez-Martin E, Scott KA, van Schalkwyk OL et al: Detection of Pathogen Exposure in African Buffalo Using Non-Specific Markers of Inflammation. Frontiers in immunology 2017, 8:1944.

20. Bronsvoort BM, Parida S, Handel I, McFarland S, Fleming L, Hamblin P, Kock R: Serological survey for foot-and-mouth disease virus in wildlife in eastern Africa and estimation of test parameters of a nonstructural protein enzyme-linked immunosorbent assay for buffalo. Clin Vaccine Immunol 2008, 15(6):1003–1011.

21. Di Nardo A, Libeau G, Chardonnet B, Chardonnet P, Kock RA, Parekh K, Hamblin P, Li Y, Parida S, Sumption KJ: Serological profile of foot-and-mouth disease in wildlife populations of West and Central Africa with special reference to Syncerus caffer subspecies. Veterinary research 2015, 46(1):77.

22. Brückner GK, Vosloo W, Du Plessis BJ, Kloeck PE, Connoway L, Ekron MD, Weaver DB, Dickason CJ, Schreuder FJ, Marais T et al: Foot and mouth disease: the experience of South Africa. Revue scientifique et technique (International Office of Epizootics) 2002, 21(3):751–764.

23. Weaver GV, Domenech J, Thiermann AR, Karesh WB: FOOT AND MOUTH DISEASE: A LOOK FROM THE WILD SIDE. Journal of Wildlife Diseases 2013, 49(4):759–785, 727.

24. Bengis RG, Thomson GR, Hedger RS, De Vos V, Pini A: Foot-and-mouth disease and the African buffalo (Syncerus caffer). 1. Carriers as a source of infection for cattle. The Onderstepoort journal of veterinary research 1986, 53(2):69–73.

25. Condy JB, Hedger RS, Hamblin C, Barnett IT: The duration of the foot-and-mouth disease virus carrier state in African buffalo (i) in the individual animal and (ii) in a free-living herd. Comparative immunology, microbiology and infectious diseases 1985, 8(3-4):259–265.

26. Fukase E: The initial cost estimate of the global FAO/OIE strategy for the contorl of foot and mouth disease. In: The Global Foot and Mouth Disease Control Strategy Strengthening animal health systems through improved control of major diseases 2012.

27. Omondi G, Alkhamis MA, Obanda V, Gakuya F, Sangula A, Pauszek S, Perez A, Ngulu S, van Aardt R, Arzt J et al: Phylogeographical and cross-species transmission dynamics of SAT1 and SAT2 foot-and-mouth disease virus in Eastern Africa. Mol Ecol 2019, 28(11):2903–2916.

28. Jori F, Etter E: Transmission of foot and mouth disease at the wildlife/livestock interface of the Kruger National Park, South Africa: Can the risk be mitigated? Preventive veterinary medicine 2016, 126:19–29.

29. Bastos ADS, Haydon DT, Sangaré O, Boshoff CI, Edrich JL, Thomson GR: The implications of virus diversity within the SAT 2 serotype for control of foot-and-mouth disease in sub-Saharan Africa. The Journal of general virology 2003, 84(Pt 6):1595–1606.

30. Hall MD, Knowles NJ, Wadsworth J, Rambaut A, Woolhouse ME: Reconstructing geographical movements and host species transitions of foot-and-mouth disease virus serotype SAT 2. mBio 2013, 4(5):e00591–00513.

31. Dyason E: Summary of foot-and-mouth disease outbreaks reported in and around the Kruger National Park, South Africa, between 1970 and 2009. J S Afr Vet Assoc 2010, 81(4):201–206.

32. Zhang ZD, Kitching RP: The localization of persistent foot and mouth disease virus in the epithelial cells of the soft palate and pharynx. Journal of comparative pathology 2001, 124(2-3):89–94.

33. Brehm KE, Ferris NP, Lenk M, Riebe R, Haas B: Highly sensitive fetal goat tongue cell line for detection and isolation of foot-and-mouth disease virus. Journal of clinical microbiology 2009, 47(10):3156–3160.

34. Fray MD, Supple EA, Morrison WI, Charleston B: Germinal centre localization of bovine viral diarrhoea virus in persistently infected animals. The Journal of general virology 2000, 81(Pt 7):1669–1673.

35. Botha A, Lease HM, Fuller A, Mitchell D, Hetem RS: Biologging subcutaneous temperatures to detect orientation to solar radiation remotely in savanna antelope. J Exp Zool A Ecol Integr Physiol 2019, 331(5):267–279.

36. Callahan JD, Brown F, Osorio FA, Sur JH, Kramer E, Long GW, Lubroth J, Ellis SJ, Shoulars KS, Gaffney KL et al: Use of a portable real-time reverse transcriptase-polymerase chain reaction assay for rapid detection of foot-and-mouth disease virus. Journal of the American Veterinary Medical Association 2002, 220(11):1636–1642.

37. Colenutt C, Brown E, Nelson N, Paton DJ, Eblé P, Dekker A, Gonzales JL, Gubbins S: Quantifying the Transmission of Foot-and-Mouth Disease Virus in Cattle via a Contaminated Environment. mBio 2020, 11(4).

38. OIE: OIE Terrestrial Manual, Chapter 3.1.8 Foot and mouth disease. In.; 2021.

39. Barnett PV, Statham RJ, Vosloo W, Haydon DT: Foot-and-mouth disease vaccine potency testing: determination and statistical validation of a model using a serological approach. Vaccine 2003, 21(23):3240–3248.

40. Reid E, Juleff N, Windsor M, Gubbins S, Roberts L, Morgan S, Meyers G, Perez-Martin E, Tchilian E, Charleston B et al: Type I and III IFNs Produced by Plasmacytoid Dendritic Cells in Response to a Member of the Flaviviridae Suppress Cellular Immune Responses. Journal of immunology (Baltimore, Md : 1950) 2016, 196(10):4214–4226.

41. Vosloo W, Bastos AD, Kirkbride E, Esterhuysen JJ, van Rensburg DJ, Bengis RG, Keet DW, Thomson GR: Persistent infection of African buffalo (Syncerus caffer) with SAT-type foot-and-mouth disease viruses: rate of fixation of mutations, antigenic change and interspecies transmission. The Journal of general virology 1996, 77 (Pt 7):1457–1467.

42. Ferris NP, Condy JB, Barnett IT, Armstrong RM: Experimental infection of eland (Taurotrages oryx), sable antelope (Ozanna grandicomis) and buffalo (Syncerus caffer) with foot-and-mouth disease virus. Journal of comparative pathology 1989, 101(3):307–316.

43. Ramulongo TD, Maree FF, Scott K, Opperman P, Mutowembwa P, Theron J: Pathogenesis, biophysical stability and phenotypic variance of SAT2 foot-and-mouth disease virus. Veterinary microbiology 2020, 243:108614.

44. Thomson GR, Vosloo W, Esterhuysen JJ, Bengis RG: Maintenance of foot and mouth disease viruses in buffalo (Syncerus caffer Sparrman, 1779) in southern Africa. Revue scientifique et technique (International Office of Epizootics) 1992, 11(4):1097–1107.

45. Gainaru MD, Thomson GR, Bengis RG, Esterhuysen JJ, Bruce W, Pini A: Foot-and-mouth disease and the African buffalo (Syncerus caffer). II. Virus excretion and transmission during acute infection. The Onderstepoort journal of veterinary research 1986, 53(2):75–85.

46. Giorgakoudi K, Gubbins S, Ward J, Juleff N, Zhang Z, Schley D: Using Mathematical Modelling to Explore Hypotheses about the Role of Bovine Epithelium Structure in Foot-And-Mouth Disease Virus-Induced Cell Lysis. PloS one 2015, 10(10):e0138571.

47. Cortey M, Ferretti L, Pérez-Martín E, Zhang F, de Klerk-Lorist L-M, Scott K, Freimanis G, Seago J, Ribeca P, van Schalkwyk L et al: Persistent Infection of African Buffalo (Syncerus caffer) with Foot-and-Mouth Disease Virus: Limited Viral Evolution and No Evidence of Antibody Neutralization Escape. Journal of virology 2019, 93(15):e00563–00519.

48. Ferretti L, Perez-Martin E, Zhang F, de Klerk-Lorist LM, Van Schalkwyk L, Maree F, Charleston B, Ribeca P: Pervasive within-host recombination and epistasis as major determinants of the molecular evolution of the Foot-and Mouth Disease Virus capsidBioRxiv 2018.

