## Additional material for "Viral dynamics and immune responses to foot-and-mouth disease virus in African buffalo *(Syncerus caffer)*"

**Additional figure 1**. Normal body temperature in African buffalo. (A) The black line is the fitted nonlinear curve, while the green points represent the data from 11 animals, with temperatures collected every 5 minutes. (B) Residuals over time from the nonlinear regression in A. (C) A scatter plot showing the range of residuals which were found to vary between -1.057 and 1.042, which we assume is normal physiological variation.

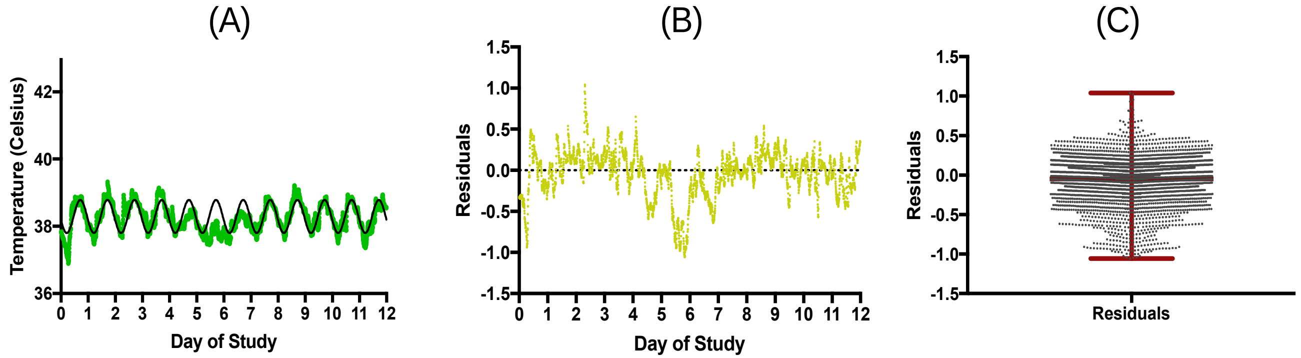

**Additional figure 2**. White blood cell (WBC) count from all needle infected (NI) and contact (Co) animals from day 0 to day 28 post virus exposure. e study

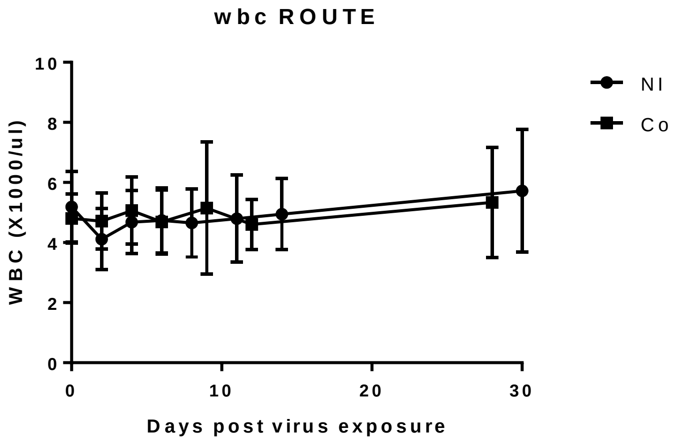

**Additional figure 3.** Boxplots showing PCR values (FMDV GCN/ml) stratified by virus isolation category (negative/positive) in probang and tonsil swab.

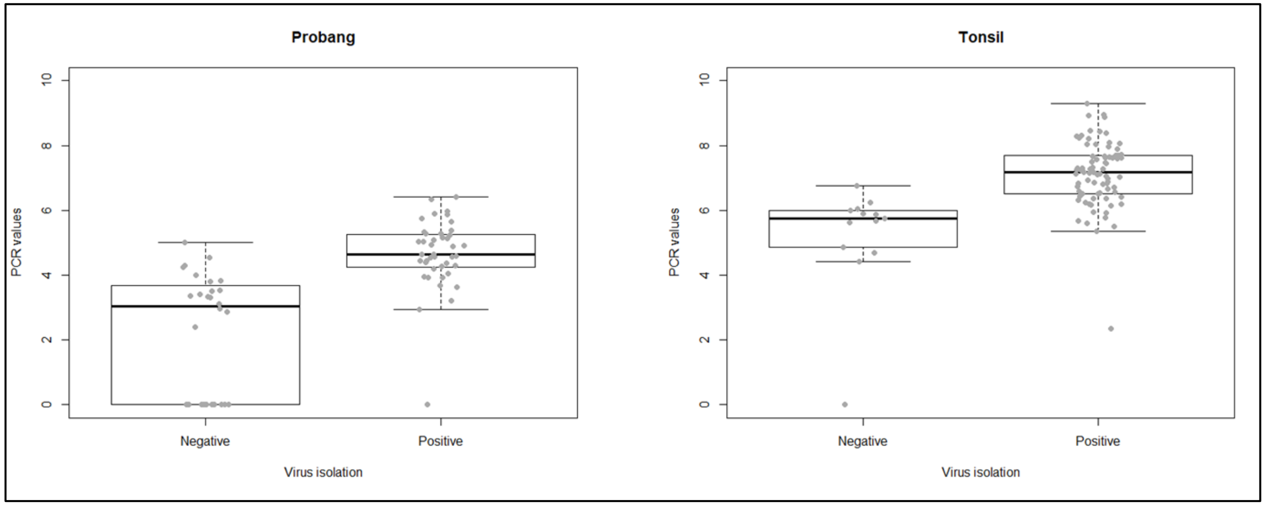

**Supplementary table 1** Median values (minimum-maximum) and the Kruskal-Wallis statistics of virus load, serology and hematology values stratified by serotype (SAT1, SAT2 and SAT3).

| Parameter | Needle infected -Median (min-max) | | | P value | Contact - Median (min-max) | | | P value |
|---|---|---|---|---|---|---|---|---|
| | **Sat1** | **Sat2** | **Sat3** | | **Sat1** | **Sat2** | **Sat3** | |
| ***VIROLOGY*** | | | | | | | | |
| **Virus load in serum** | | | | | | | | |
| •AUC ($log_{10}$) | 3.37 (3.19-3.61) | 3.48 (3.02-3.53) | 3.01 (2.92-3.08) | 0.058 | 3.42 (3.36-3.53) | 2.57 (2.00-3.15) | 3.24 (3.01-3.41) | **0.028** |
| •Day viremia starts | 2 (2-2) | 2 (2-2) | 2 (2-2) | - | 2 (2-2) | 4 (2-4) | 5 (4-12) | **0.017** |
| •Day viremia peaks | 2 (2-4) | 2 (2-2) | 2 (2-2) | 0.368 | 4 (4-4) | 4 (4-6) | 6 (4-9) | 0.150 |
| •Peak value | 7.07 (6.31-7.67) | 6.25 (5.20-7.31) | 5.58 (5.22-7.46) | 0.232 | 5.71 (5.51-8.39) | 4.13 (2.50-5.64) | 6.2 (5.59-6.22) | 0.100 |
| •Duration (days) | 5 (4-6) | 6 (4-6) | 4 (4-4) | 0.111 | 6.5 (6.5-6.5) | 4.5 (2.5-6.5) | 2.5 (2.5-2.5) | **0.021** |
| **Virus load in tonsils** | | | | | | | | |
| •AUC ($log_{10}$) | 5.43 (5.36-5.49) | 5.39 (5.26-5.42) | 5.27 (5.11-5.36) | **0.048** | 5.27 (4.81-5.37) | 5.16 (5.08-5.27) | 5.14 (5.12-5.15) | 0.523 |
| •First day detected | 2 (2-2) | 2 (2-2) | 2 (2-4) | 0.368 | 2 (2 - 2) | 3 (2- 4) | 5 (2- 12) | 0.088 |
| •Day peaks | 3 (2-4) | 5 (4-6) | 6 (4-6) | **0.050** | 6 (4-6) | 4 (4-9) | 12 (9-12) | **0.045** |
| •Peak value | 9.54 (9.12-9.98) | 9.43 (9.32-9.63) | 8.67 (7.99-9.70) | 0.232 | 9.13 (7.93-9.63) | 9.13 (8.46-9.99) | 7.70 (7.60-8.29) | **0.076** |
| **Nasal swab** | | | | | | | | |
| •First day detected | 3 (2-6) | 6.5 (2-11) | 2 (2-11) | 0.820 | 4 (4-4) | 4 (2-9) | 9 (9-9) | 0.142 |
| •Day peaks | 4 (2-6) | 6.5 (2-11) | 11 (2 -11) | 0.698 | 6.5 (4-9) | 9 (2-9) | 9 (9-9) | 0.498 |
| •Peak value | 3.96 (2.78-6.57) | 2.95 (2.53-5.29) | 3.83 (0-4.95) | 0.618 | 5.33 (4.20-6.47) | 3.44 (2.42-4.16) | 3.17 (2.74-3.24) | 0.119 |

| | | | | | | | | |
|---|---|---|---|---|---|---|---|---|
| **Day virus first detected** | 2 (2-2) | 2 (2-2) | 2 (2-2) | - | 2 (2-2) | 3 (2-4) | 4 (2-9) | 0.103 |
| ***SEROLOGY*** | | | | | | | | |
| **VNT** | | | | | | | | |
| •First day positive | 4 (2 -6) | 6 (6-6) | 6 (2-6) | 0.134 | 4 (2-6) | 9 (6-12) | 9 (6-12) | **0.021** |
| •First Day protective titre | 6 (4-6) | 6 (6-8) | 6 (6-8) | 0.294 | 6 (6-9) | 10.5 (9-12) | 9 (9-12) | **0.050** |
| •Day peaks | 22 (14-30) | 11 (11-14) | 12.5 (8-14) | 0.063 | 12 (9-12) | 12 (12-28) | 12 (12-28) | 0.287 |
| •Peak value ($log_{10}$) | 3.15 (3.15-3.15) | 3.15 (3.15-3.15) | 3.15 (3.15-3.15) | - | 3.15 (3.15-3.15) | 3.15 (3.01-3.15) | 2.78 (2.25-3.15) | 0.070 |
| •Response time | 2 (0-4) | 4 (4-4) | 4 (0-4) | 0.134 | 2 (0-4) | 6 (4-8) | 7 (2-8) | 0.106 |
| **NSP** | | | | | | | | |
| •First day positive | 8 (6-8) | 8 (6-11) | 8 (6-8) | 0.829 | 10.5 (9-12) | 10.5 (9-12) | 12 (12 – 28) | 0.187 |
| •Response time | 6 (4-6) | 6 (4-9) | 6 (4-6) | 0.829 | 8.5 (7-10) | 7.5 (5-10) | 10 (8-24) | 0.287 |
| **Interferon γ** | | | | | | | | |
| •First day detected | 2 (2-2) | 2 (2-2) | 2 (2-2) | - | 2 (2-2) | 2 (2-2) | 2 (2-2) | - |
| •AUC ($log_{10}$) | 3.71 (3.17-3.99) | 3.90 (3.84-4.18) | 3.73 (3.47-4.45) | 0.276 | 3.69 (6.66-3.76) | 3.83 (3.51-4.49) | 3.75 (3.74-3.85) | 0.406 |
| •Day peak | 5 (2-6) | 4 (2-8) | 4 (2-14) | 0.995 | 4 (4-4) | 4 (2-4) | 4(4-6) | 0.233 |
| •Peak value (µg/ml) | 4.97 (3.91-5.48) | 5.09 (4.04-11.47) | 4.48 (4.04-5.22) | 0.608 | 6.52 (6.13-7.86) | 6.95 (3.67-23.09) | 4.98 (4.72-5.81) | 0.176 |
| •Response time | 0 (0-0) | 0 (0-0) | 0 (0-0) | - | 0 (0-0) | 0 (-2- 0) | -2 (-2 - 0) | 0.364 |
| **Type I/III IFN** | | | | | | | | |
| •AUC ($log_{10}$) | 2.65 (1.96-2.79) | 2.69 (2.47-3.02) | 2.15 (1.49-2.61) | 0.219 | 2.63 (2.14-2.82) | 2.42 (2.32-2.51) | 1.77 (1.73-2.66) | 0.356 |
| •Day peak | 2 (2-4) | 2 (2-2) | 4 (2-6) | 0.238 | 5 (4-6) | 6 (4-6) | 4 (0-6) | 0.435 |
| •Peak value (iu/ml) | 3.17 (1.24-3.65) | 2.93 (2.54-5.24) | 1.41 (0.66-3.83) | 0.397 | 3.03 (1.38-5.07) | 2.27 (1.79-2.90) | 1.97 (1.88-2.44) | 0.749 |
| •Response time | 0 (0-0) | 0 (0-0) | 0 (0-0) | - | 2 (0-2) | 0 (-2- 0) | 1 (0 2) | 0.094 |
| ***HEMATOTOGY*** | | | | | | | | |
| **HAPTOGLOBULIN** | | | | | | | | |
| •First day positive | 2 (2-2) | 2 (2-2) | 2 (2-2) | **-** | 4 (2-4) | 4 (2-6) | 4 (2-12) | 0.903 |
| •AUC ($log_{10}$) | 15.53 (15.46-15.58) | 16.01 (15.99-16.06) | 15.23 (14.66-15.66) | **0.018** | 15.90 (15.47-16.12) | 15.71 (15.40-16.0) | 13.13 (10.21-15.64) | 0.138 |
| •Day peaks | 8 (4-11) | 11 (2-11) | 6 (4-11) | 0.621 | 9 (9-12) | 12 (6-12) | 9 (6-9) | 0.294 |
| •Peak value (ng/ml) | 641082 (509902-671178) | 666807 (643331-687700) | 596071 (293986-660863) | 0.174 | 654360 (537771-671676) | 629915 (560970-659085) | 154080 (3784-635514) | 0.161 |
| •Response time | 0 (0-0) | 0 (0-0) | 0 (0-0) | - | 2 (0-2) | 2 (-2-2) | 0.5 (-2 - 4) | 0.962 |
| **SAA** | | | | | | | | |
| •First day detected | 2 (2-2) | 2 (2-2) | 2 (2-2) | - | 2 (2-2) | 3 (2-6) | 2 (2-2) | - |
| •AUC ($log_{10}$) | 11.35 (11.12-11.46) | 11.56 (10.83-11.75) | 11.19 (10.95-11.49) | 0.499 | 11.46 (10.90-12.61) | 11.25 (11.04-11.72) | 11.72 (11.30-11.74) | 0.446 |
| •Day peaks | 4 (2-6) | 4 (4-6) | 5 (2-6) | 0.852 | 6 (4-28) | 6 (6-6) | 6 (6-9) | 0.732 |
| •Peak value (ng/ml) | 12053 (10974-12822) | 12364 (7711-15000) | 12044 (10792-13825) | 0.926 | 14532 (9646-14883) | 13974 (13159-15000) | 14536 (13936-15000) | 0.722 |
| •Response time | 0 (0-0) | 0 (0-0) | 0 (0-0) | - | 0 (0-0) | 1 (-2 - 2) | -2 (-2 0) | 0.219 |

**Supplementary table 2** Median values (minimum - maximum) and the Kruskal-Wallis statistics of virus load, serology and hematology values stratified by method of infection (needle infected vs contact)

| Parameter | Median (min-max) | | P value |
|---|---|---|---|
| | Needle infected | Contact | |
| ***VIROLOGY*** | | | |
| **Virus load in serum** | | | |
| •AUC ($log_{10}$) | 3.22 (2.92-3.61) | 3.24 (2.00-3.53) | 0.389 |
| •Day viremia starts | 2 (2-2) | 2 (2-12) | **0.002** |
| •Day viremia peaks | 2 (2-4) | 4 (4-9) | **<0.001** |
| •Peak value | 6.35 (5.20-7.67) | 5.59 (2.50-8.39) | 0.085 |
| •Duration (days) | 4 (4-6) | 4.5 (2.5-6.5) | 0.613 |
| **Virus load in tonsils** | | | |
| •AUC ($log_{10}$) | 5.36 (5.11-5.49) | 5.15 (4.82-5.37) | **0.004** |
| •First day detected | 2 (2-2) | 4 (2 – 12) | 0.057 |
| •Day peaks | 4 (2-6) | 6 (4-12) | 0.064 |
| •Peak value | 9.34 (7.99-9.98) | 8.92 (7.60-9.99) | 0.103 |
| **Nasal swab** | | | |
| •First day detected | 2 (2-11) | 6.5 (2-9) | 0.369 |
| •Day peaks | 4 (2-11) | 9 (2-9) | 0.670 |
| •Peak value | 3.42 (0-6.57) | 3.34 (2.42-6.47) | 0.728 |
| **Day virus first detected** | 2 (2-2) | 2 (2-9) | 0.014 |
| ***SEROLOGY*** | | | |
| **VNT** | | | |
| •First day positive | 4 (2-4) | 6 (2 – 12) | 0.062 |
| •First Day protective titre | 6 (4-8) | 9 (6-12) | **0.002** |
| •Day peaks | 14 (8-30) | 12 (9-28) | 0.751 |
| •Peak value (log10) | 3.15 (3.15-3.15) | 3.15 (2.25-3.15) | **0.033** |
| •Response time | 4 (0-4) | 4 (0-8) | 0.132 |
| **NSP** | | | |
| •First day positive | 8 (6-11) | 12 (9-28) | **<0.001** |
| •Response time | 6 (4-9) | 8 (5-24) | **0.001** |
| **Interferon γ** | | | |
| •First day detected | 2 (2-2) | 2 (2-2) | - |
| •AUC ($log_{10}$) | 3.83 (3.17-4.45) | 3.74 (3.51-4.49) | 0.623 |
| •Day peak | 5 (2-14) | 4 (2-6) | 0.696 |
| •Peak value (µg/ml) | 4.90 (3.91-11.47) | 6.13 (3.67-23.09) | **0.036** |
| •Response time | 0 (0-0) | 0 (-2 0) | **0.031** |
| • Duration (days) | 14 (6-14) | 9 (6-14) | 0.052 |
| **Type I/III IFN** | | | |
| •AUC ($log_{10}$) | 2.58 (1.49-3.02) | 2.47 (1.73-2.82) | 0.355 |
| •Day peak | 2 (2-6) | 6 (0-6) | **0.013** |
| •Peak value (iu/ml) | 2.93 (0.66-5.24) | 2.18 (1.38-5.07) | 0.580 |
| •Response time | 0 (0-0) | 0 (-2 2) | 0.152 |
| •Duration (days) | 6 (6-6) | 6 (0-9) | 0.528 |
| ***HEMATOTOGY*** | | | |
| **HAPTOGLOBULIN** | | | |
| •First day positive | 2 (2-2) | 4 (2 – 12) | **<0.001** |
| •AUC ($log_{10}$) | 15.56 (14.66-16.06) | 15.64 (10.21-16.12) | 0.902 |
| •Day peaks | 8 (2-11) | 9 (6-12) | 0.118 |
| •Peak value (ng/ml) | 644713 (293986-687700) | 635514 (3784-671676) | 0.325 |
| •Response time | 0(0-0) | 2 (-2-4) | 0.057 |
| **SAA** | | | |

| | | | |
|---|---|---|---|
| •First day detected | 2(2-2) | 2 (2-6) | - |
| •AUC ($log_{10}$) | 11.35 (10.83-11.75) | 11.38 (10.90-12.61) | 0.460 |
| •Day peaks | 4 (2-6) | 6 (4-28) | **0.004** |
| •Peak value (ng/ml) | 12154 (7711-15000) | 14217(9646-15000) | **0.007** |
| •Response time | 0 (0-0) | 0 (-2 2) | 0.622 |
| •Duration (days) | 11 (8-14) | 12 (9-28) | **0.002** |
